## Supplementary figures 1-4 and supplementary tables 1-3 for "Unbiased placental secretome characterization identifies candidates for pregnancy complications"

S1. Gene Enrichment for non-secreted proteins detected in mouse placental endocrine (Jz +Tpbpa sorted cells)

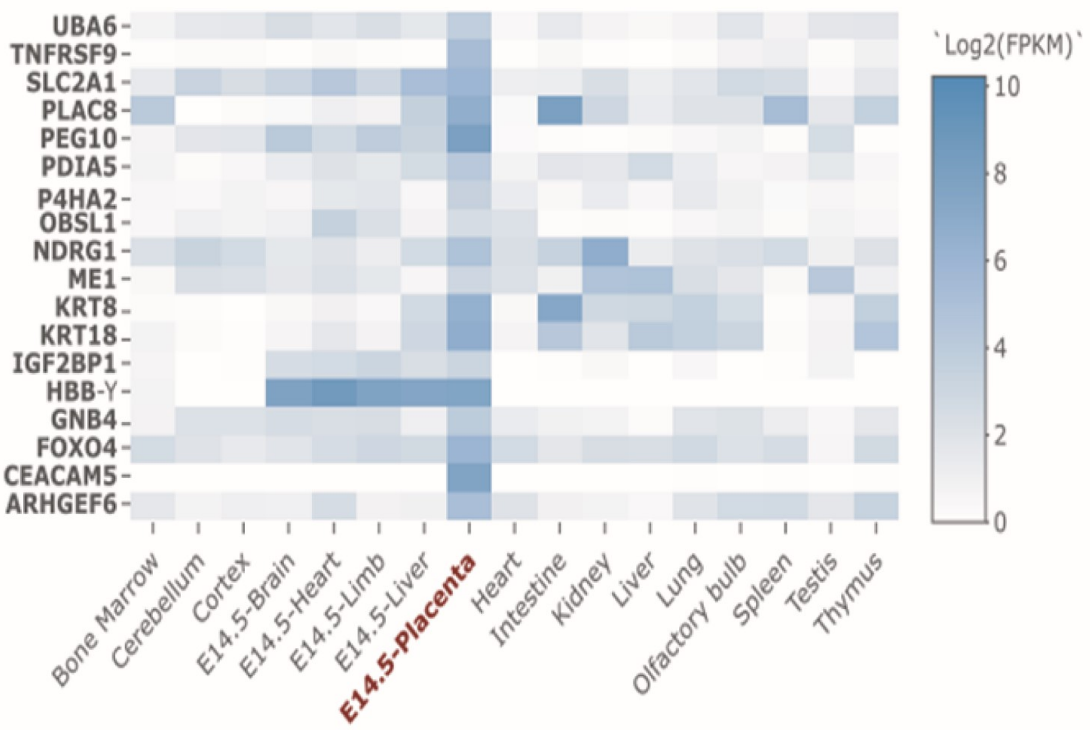


Mouse specific enrichment analysis ( <10 fold) for the non-secreted proteins. 18 non secreted proteins are enriched in the mouse placenta

S2: Analysis of the 31 secreted proteins reported to be expressed by the mouse but not human placenta

A.

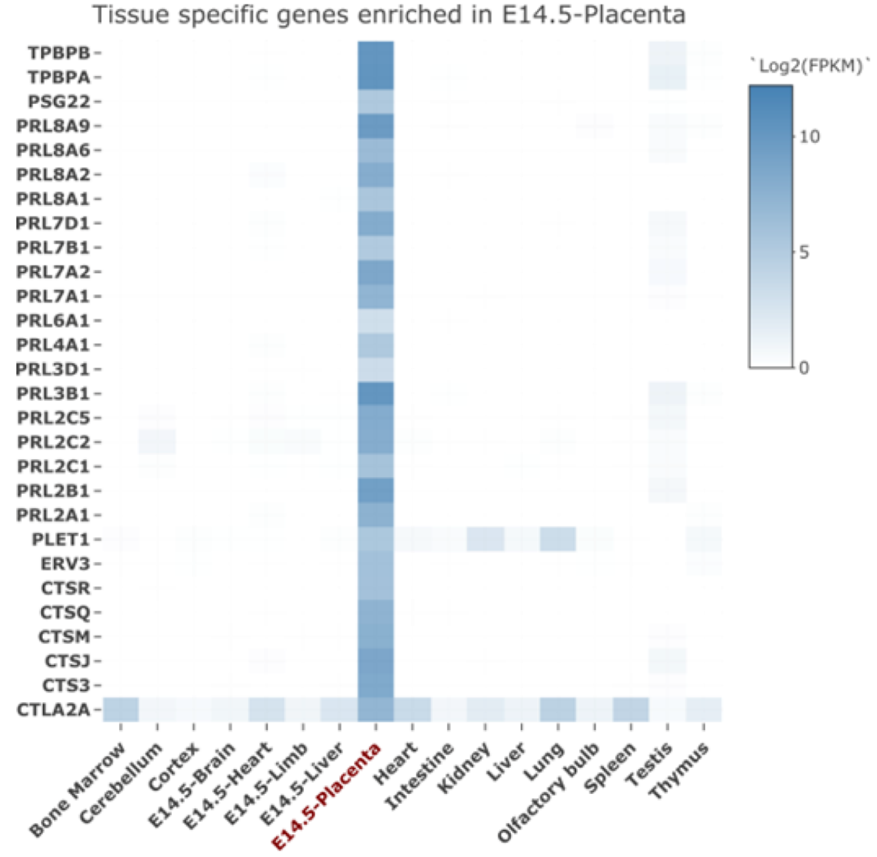


Mouse specific enrichment analysis ( <10 fold) for the 31 secreted proteins specific to the mouse (28 were detected to be enriched in the placenta).

B.

Biological Process

| #Term ID | Term description | # obs. | FDR |
|---|---|---|---|
| *GO:0010469* | Regulation of signaling receptor activity | 14 | 6.45E-13 |
| *GO:0065009* | Regulation of molecular function | 15 | 0.00023 |
| *GO:0048583* | Regulation of response to stimulus | 15 | 0.00071 |

Molecular Function

| #Term ID | Term description | # obs. | FDR |
|---|---|---|---|
| *GO:0005179* | Hormone activity | 14 | 1.39E-22 |
| *GO:0098772* | Molecular function regulator | 15 | 1.68E-09 |
| *GO:0008234* | Cysteine-type peptidase activity | 5 | 1.48E-05 |
| *GO:0004197* | Cysteine-type endopeptidase activity | 3 | 0.00073 |
| *GO:0005515* | Protein binding | 16 | 0.0107 |

Protein Domains

| #Term ID | Term description | # obs. | FDR |
|---|---|---|---|
| MMU-8939242 | RUNX1 regulates transcription of genes involved in differentiation of keratinocytes | 4 | 1.07E-07 |
| MMU-1679131 | Trafficking and processing of endosomal TLR | 4 | 2.83E-07 |
| MMU-1442490 | Collagen degradation | 4 | 1.04E-05 |
| MMU-2022090 | Assembly of collagen fibrils and other multimeric structures | 4 | 1.04E-05 |
| MMU-2132295 | MHC class II antigen presentation | 21 | 4.99E-08 |

S3. Expression of ANGPT2, MIF and IGF2 at the maternal-fetal interface in early human pregnancy

A. ANGPT2 expression

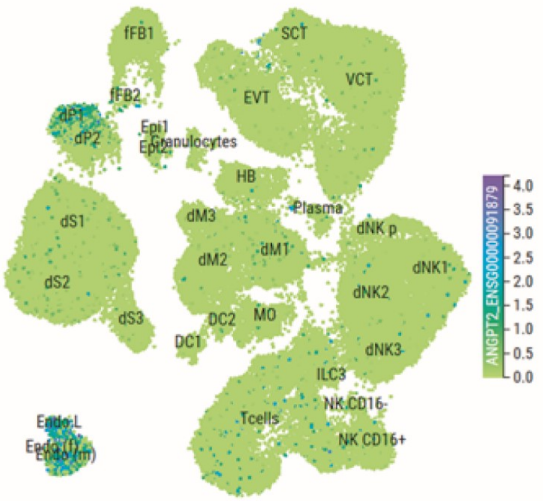


B. MIF expression

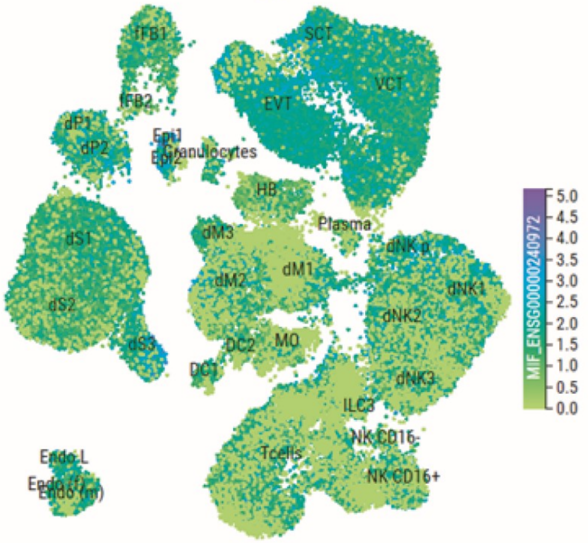


C. IGF2 expression

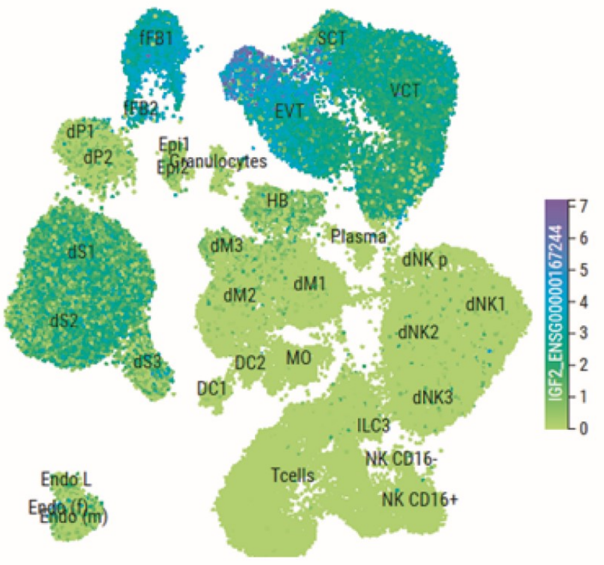

S4. Concentrations of placenta proteins in human pregnancy samples

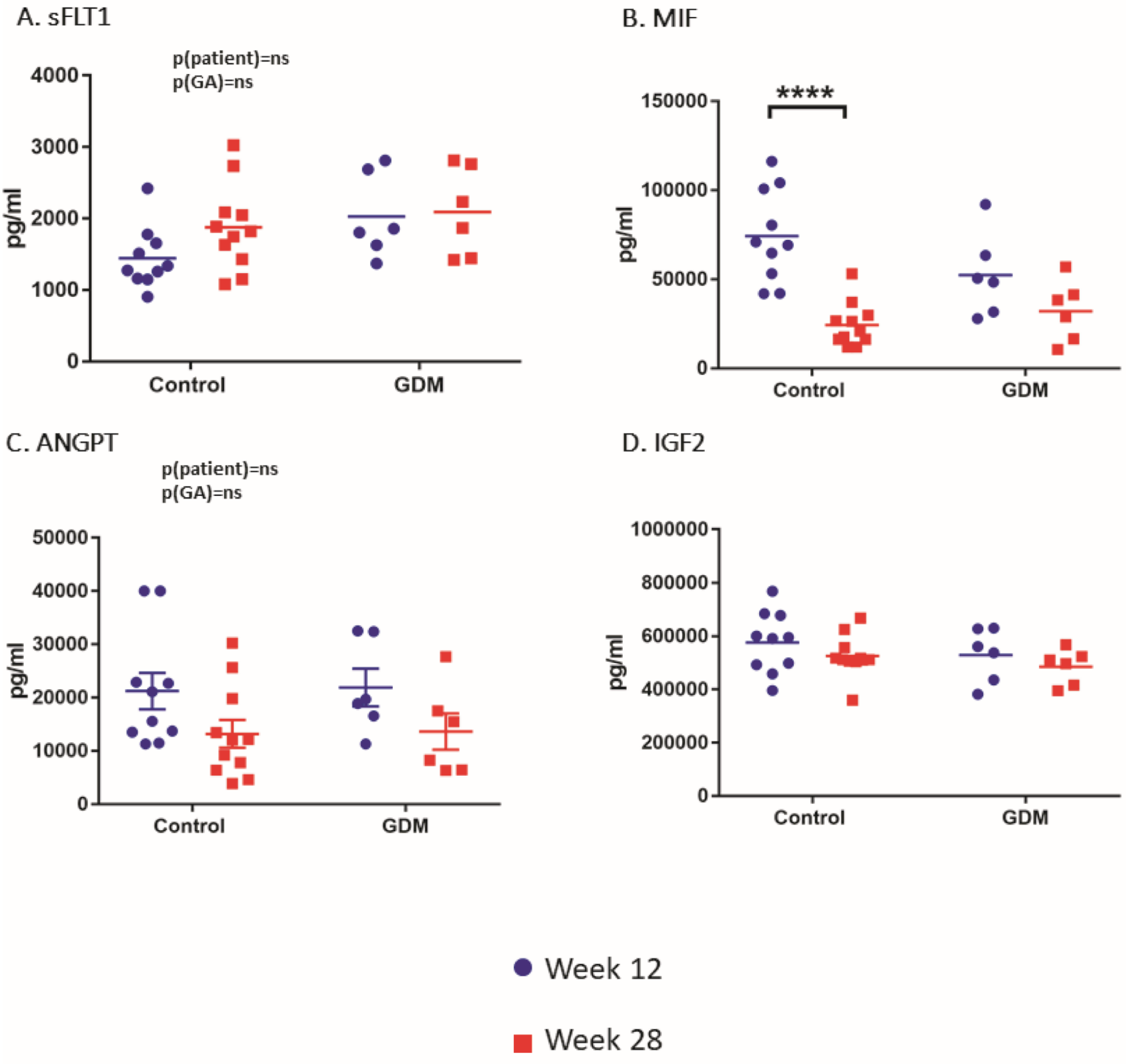

### S1 A. Panther pathway analysis for non secreted proteins from the Jz

| PANTHER Protein Class | Non secreted protein list | Fold Enrichment | P-value | FDR |
|---|---|---|---|---|
| chaperonin (PC00073) | 9 | 27.43 | 1.17E-09 | 2.53E-08 |
| ribosomal protein (PC00202) | 53 | 13.21 | 5.06E-38 | 4.93E-36 |
| translation initiation factor (PC00224) | 18 | 11.89 | 4.56E-13 | 1.48E-11 |
| translational protein (PC00263) | 86 | 11.83 | 2.41E-58 | 4.70E-56 |
| aminoacyl-tRNA synthetase (PC00047) | 9 | 10.49 | 8.34E-07 | 1.16E-05 |
| translation factor (PC00223) | 24 | 10.01 | 1.49E-15 | 5.82E-14 |
| tubulin (PC00228) | 5 | 9.9 | 3.14E-04 | 2.66E-03 |
| peroxidase (PC00180) | 5 | 8.61 | 5.46E-04 | 3.80E-03 |
| chaperone (PC00072) | 16 | 8.57 | 5.89E-10 | 1.44E-08 |
| vesicle coat protein (PC00235) | 10 | 8.09 | 1.61E-06 | 2.09E-05 |
| actin and actin related protein (PC00039) | 6 | 7.92 | 2.25E-04 | 2.00E-03 |
| translation elongation factor (PC00222) | 3 | 7.92 | 9.20E-03 | 4.60E-02 |
| G-protein (PC00020) | 18 | 5.17 | 6.12E-08 | 9.95E-07 |
| histone (PC00118) | 8 | 4.8 | 4.65E-04 | 3.49E-03 |
| small GTPase (PC00208) | 8 | 4.73 | 5.09E-04 | 3.68E-03 |
| heterotrimeric G-protein (PC00117) | 6 | 4.57 | 2.97E-03 | 1.70E-02 |
| actin or actin-binding cytoskeletal protein ( | 26 | 4.36 | 1.95E-09 | 3.80E-08 |
| ligase (PC00142) | 6 | 3.96 | 5.65E-03 | 3.06E-02 |
| non-motor actin binding protein (PC00165) | 9 | 3.83 | 9.38E-04 | 5.90E-03 |
| dehydrogenase (PC00092) | 10 | 3.33 | 1.36E-03 | 8.01E-03 |
| cytoskeletal protein (PC00085) | 40 | 3.22 | 4.57E-10 | 1.27E-08 |
| microtubule or microtubule-binding cytosk | 12 | 2.46 | 5.02E-03 | 2.79E-02 |
| protease (PC00190) | 34 | 2.26 | 2.10E-05 | 2.28E-04 |
| oxidoreductase (PC00176) | 23 | 2.18 | 8.38E-04 | 5.45E-03 |
| membrane traffic protein (PC00150) | 19 | 1.95 | 8.42E-03 | 4.32E-02 |
| transferase (PC00220) | 23 | 1.85 | 5.92E-03 | 3.12E-02 |
| metabolite interconversion enzyme (PC002 | 59 | 1.64 | 3.49E-04 | 2.72E-03 |
| protein modifying enzyme (PC00260) | 49 | 1.61 | 1.35E-03 | 8.26E-03 |
| protein class (PC00000) | 363 | 1.45 | 4.27E-21 | 2.08E-19 |

**S1 B. Reactome pathway analysis of the non secreted proteins from the Jz**

| | |
|---|---|
| Analysis Type: | PANTHER Overrepresentation Test (Released 20200407) |
| Annotation Version and Release Date: | Reactome version 65 Released 2019-12-22 |
| Analyzed List: | upload_1 (Mus musculus) |
| Reference List: | Mus musculus (all genes in database) |
| Test Type: | FISHER |
| Correction: | FDR |

| Reactome pathways | Mus musculus - REFLIST (22265) | upload_1 (562) | upload_1 (expected) | upload_1 (over/under) | upload_1 (fold Enrichment) | upload_1 (raw P-value) | upload_1 (FDR) |
|---|---|---|---|---|---|---|---|
| Association of TriC/CCT with target proteins during biosynthesis (R-MMU-390471) | 10 | 8 | 0.25 | + | 31.69 | 4.52E-09 | 4.63E-08 |
| Eukaryotic Translation Elongation (R-MMU-156842) | 5 | 4 | 0.13 | + | 31.69 | 4.15E-05 | 3.17E-04 |
| Formation of the ternary complex, and subsequently, the 43S complex (R-MMU-72695) | 52 | 41 | 1.31 | + | 31.24 | 3.40E-41 | 2.55E-39 |
| Ribosomal scanning and start codon recognition (R-MMU-72702) | 59 | 46 | 1.49 | + | 30.89 | 5.93E-46 | 6.98E-44 |
| Translation initiation complex formation (R-MMU-72649) | 59 | 46 | 1.49 | + | 30.89 | 5.93E-46 | 6.52E-44 |
| Activation of the mRNA upon binding of the cap-binding complex and eIFs, and subsequent binding to 43S (R-MMU-72662) | 60 | 46 | 1.51 | + | 30.37 | 1.02E-45 | 1.06E-43 |
| PERK regulates gene expression (R-MMU-381042) | 4 | 3 | 0.1 | + | 29.71 | 4.83E-04 | 3.23E-03 |
| GTP hydrolysis and joining of the 60S ribosomal subunit (R-MMU-72706) | 110 | 82 | 2.78 | + | 29.53 | 1.29E-80 | 1.07E-77 |
| L13a-mediated translational silencing of Ceruloplasmin expression (R-MMU-156827) | 109 | 81 | 2.75 | + | 29.44 | 1.53E-79 | 5.05E-77 |
| Formation of a pool of free 40S subunits (R-MMU-72689) | 99 | 73 | 2.5 | + | 29.21 | 1.89E-71 | 5.18E-69 |
| Cap-dependent Translation Initiation (R-MMU-72737) | 117 | 83 | 2.95 | + | 28.1 | 2.53E-80 | 1.39E-77 |
| Eukaryotic Translation Initiation (R-MMU-72613) | 117 | 83 | 2.95 | + | 28.1 | 2.53E-80 | 1.04E-77 |
| Nonsense Mediated Decay (NMD) independent of the Exon Junction Complex (EJC) (R-MMU-975956) | 92 | 65 | 2.32 | + | 27.99 | 9.74E-63 | 1.79E-60 |
| SRP-dependent cotranslational protein targeting to membrane (R-MMU-1799339) | 90 | 62 | 2.27 | + | 27.29 | 2.35E-59 | 2.98E-57 |
| Regulation of RUNX2 expression and activity (R-MMU-8939902) | 50 | 34 | 1.26 | + | 26.94 | 8.80E-33 | 4.68E-31 |
| AUF1 (hnRNP D0) binds and destabilizes mRNA (R-MMU-450408) | 55 | 37 | 1.39 | + | 26.65 | 1.88E-35 | 1.24E-33 |
| Formation of the active cofactor, UDP-glucuronate (R-MMU-173599) | 3 | 2 | 0.08 | + | 26.41 | 5.76E-03 | 3.02E-02 |
| Cross-presentation of soluble exogenous antigens (endosomes) (R-MMU-1236978) | 48 | 32 | 1.21 | + | 26.41 | 1.01E-30 | 3.71E-29 |
| Ubiquitin-dependent degradation of Cyclin D (R-MMU-75815) | 50 | 33 | 1.26 | + | 26.15 | 1.53E-31 | 6.84E-30 |
| Autodegradation of the E3 ubiquitin ligase COP1 (R-MMU-349425) | 51 | 33 | 1.29 | + | 25.63 | 2.47E-31 | 1.07E-29 |
| p53-Independent G1/S DNA damage checkpoint (R-MMU-69613) | 51 | 33 | 1.29 | + | 25.63 | 2.47E-31 | 1.04E-29 |
| p53-Independent DNA Damage Response (R-MMU-69610) | 51 | 33 | 1.29 | + | 25.63 | 2.47E-31 | 1.02E-29 |
| Ubiquitin Mediated Degradation of Phosphorylated Cdc25A (R-MMU-69601) | 51 | 33 | 1.29 | + | 25.63 | 2.47E-31 | 9.93E-30 |
| Regulation of ornithine decarboxylase (ODC) (R-MMU-350562) | 50 | 32 | 1.26 | + | 25.36 | 2.62E-30 | 8.82E-29 |
| FBXL7 down-regulates AURKA during mitotic entry and in early mitosis (R-MMU-8854050) | 54 | 34 | 1.36 | + | 24.94 | 5.93E-32 | 3.05E-30 |
| HSF1 activation (R-MMU-3371511) | 8 | 5 | 0.2 | + | 24.76 | 9.71E-06 | 7.89E-05 |
| Regulation of RUNX3 expression and activity (R-MMU-8941858) | 53 | 33 | 1.34 | + | 24.67 | 6.26E-31 | 2.40E-29 |
| Degradation of GLI1 by the proteasome (R-MMU-5610780) | 55 | 34 | 1.39 | + | 24.49 | 9.37E-32 | 4.68E-30 |
| Degradation of AXIN (R-MMU-4641257) | 54 | 33 | 1.36 | + | 24.21 | 9.85E-31 | 3.69E-29 |
| GLI3 is processed to GLI3R by the proteasome (R-MMU-5610785) | 56 | 34 | 1.41 | + | 24.05 | 1.47E-31 | 6.94E-30 |
| Degradation of DVL (R-MMU-4641258) | 56 | 34 | 1.41 | + | 24.05 | 1.47E-31 | 6.74E-30 |
| Stabilization of p53 (R-MMU-69541) | 55 | 33 | 1.39 | + | 23.77 | 1.54E-30 | 5.41E-29 |
| Galactose catabolism (R-MMU-70370) | 5 | 3 | 0.13 | + | 23.77 | 7.58E-04 | 4.92E-03 |
| Nonsense Mediated Decay (NMD) enhanced by the Exon Junction Complex (EJC) (R-MMU-975957) | 113 | 67 | 2.85 | + | 23.49 | 6.39E-61 | 1.05E-58 |
| Nonsense-Mediated Decay (NMD) (R-MMU-927802) | 113 | 67 | 2.85 | + | 23.49 | 6.39E-61 | 9.57E-59 |
| NIK-->noncanonical NF-kB signaling (R-MMU-5676590) | 57 | 33 | 1.44 | + | 22.94 | 3.69E-30 | 1.17E-28 |
| Dectin-1 mediated noncanonical NF-kB signaling (R-MMU-5607761) | 57 | 33 | 1.44 | + | 22.94 | 3.69E-30 | 1.15E-28 |
| Metabolism of polyamines (R-MMU-351202) | 58 | 33 | 1.46 | + | 22.54 | 5.67E-30 | 1.70E-28 |
| CDT1 association with the CDC6:ORC:origin complex (R-MMU-68827) | 58 | 33 | 1.46 | + | 22.54 | 5.67E-30 | 1.67E-28 |
| Hedgehog ligand biogenesis (R-MMU-5358346) | 62 | 35 | 1.56 | + | 22.36 | 1.25E-31 | 6.04E-30 |

| | | | | | | |
|---|---|---|---|---|---|---|
| Transcriptional regulation by RUNX2 (R-MMU-8878166) | 61 | 34 | 1.54 + | 22.08 | 1.28E-30 | 4.59E-29 |
| SCF(Skp2)-mediated degradation of p27/p21 (R-MMU-187577) | 60 | 33 | 1.51 + | 21.79 | 1.31E-29 | 3.48E-28 |
| Asymmetric localization of PCP proteins (R-MMU-4608870) | 61 | 33 | 1.54 + | 21.43 | 1.97E-29 | 5.08E-28 |
| Oxygen-dependent proline hydroxylation of Hypoxia-inducible Factor Alpha (R-MMU-1234176) | 65 | 35 | 1.64 + | 21.33 | 4.31E-31 | 1.69E-29 |
| MAPK6/MAPK4 signaling (R-MMU-5687128) | 73 | 39 | 1.84 + | 21.17 | 2.02E-34 | 1.23E-32 |
| Autodegradation of Cdh1 by Cdh1:APC/C (R-MMU-174084) | 62 | 33 | 1.56 + | 21.09 | 2.95E-29 | 7.16E-28 |
| The role of GTSE1 in G2/M progression after G2 checkpoint (R-MMU-8852276) | 73 | 38 | 1.84 + | 20.62 | 3.05E-33 | 1.68E-31 |
| p53-Dependent G1/S DNA damage checkpoint (R-MMU-69580) | 64 | 33 | 1.62 + | 20.43 | 6.51E-29 | 1.51E-27 |
| p53-Dependent G1 DNA Damage Response (R-MMU-69563) | 64 | 33 | 1.62 + | 20.43 | 6.51E-29 | 1.49E-27 |
| Activation of NF-kappaB in B cells (R-MMU-1169091) | 64 | 33 | 1.62 + | 20.43 | 6.51E-29 | 1.47E-27 |
| Activation of BAD and translocation to mitochondria (R-MMU-111447) | 12 | 6 | 0.3 + | 19.81 | 3.13E-06 | 2.72E-05 |
| Axonal growth stimulation (R-MMU-209563) | 4 | 2 | 0.1 + | 19.81 | 8.50E-03 | 4.25E-02 |
| Chk1/Chk2(Cds1) mediated inactivation of Cyclin B:Cdk1 complex (R-MMU-75035) | 12 | 6 | 0.3 + | 19.81 | 3.13E-06 | 2.70E-05 |
| APC/C:Cdc20 mediated degradation of Securin (R-MMU-174154) | 66 | 33 | 1.67 + | 19.81 | 1.41E-28 | 3.09E-27 |
| G1/S DNA Damage Checkpoints (R-MMU-69615) | 66 | 33 | 1.67 + | 19.81 | 1.41E-28 | 3.05E-27 |
| Cellular response to hypoxia (R-MMU-1234174) | 70 | 35 | 1.77 + | 19.81 | 3.07E-30 | 1.01E-28 |
| RUNX1 regulates transcription of genes involved in differentiation of HSCs (R-MMU-8939236) | 66 | 33 | 1.67 + | 19.81 | 1.41E-28 | 3.01E-27 |
| Regulation of PTEN stability and activity (R-MMU-8948751) | 68 | 34 | 1.72 + | 19.81 | 2.08E-29 | 5.27E-28 |
| Regulation of RAS by GAPs (R-MMU-5658442) | 68 | 34 | 1.72 + | 19.81 | 2.08E-29 | 5.19E-28 |
| Assembly of the pre-replicative complex (R-MMU-68867) | 67 | 33 | 1.69 + | 19.51 | 2.05E-28 | 4.34E-27 |
| Orc1 removal from chromatin (R-MMU-68949) | 70 | 34 | 1.77 + | 19.24 | 4.40E-29 | 1.04E-27 |
| Regulation of mRNA stability by proteins that bind AU-rich elements (R-MMU-450531) | 86 | 41 | 2.17 + | 18.89 | 1.36E-34 | 8.65E-33 |
| Cyclin E associated events during G1/S transition (R-MMU-69202) | 70 | 33 | 1.77 + | 18.68 | 6.19E-28 | 1.25E-26 |
| CDK-mediated phosphorylation and removal of Cdc6 (R-MMU-69017) | 71 | 33 | 1.79 + | 18.41 | 8.87E-28 | 1.76E-26 |
| Cdc20:Phospho-APC/C mediated degradation of Cyclin A (R-MMU-174184) | 71 | 33 | 1.79 + | 18.41 | 8.87E-28 | 1.74E-26 |
| APC/C:Cdh1 mediated degradation of Cdc20 and other APC/C:Cdh1 targeted proteins in late mitosis/early G1 (R-MMU-174178) | 71 | 33 | 1.79 + | 18.41 | 8.87E-28 | 1.72E-26 |
| APC:Cdc20 mediated degradation of cell cycle proteins prior to satisfation of the cell cycle checkpoint (R-MMU-179419) | 72 | 33 | 1.82 + | 18.16 | 1.26E-27 | 2.42E-26 |
| Cyclin A:Cdk2-associated events at S phase entry (R-MMU-69656) | 72 | 33 | 1.82 + | 18.16 | 1.26E-27 | 2.40E-26 |
| Degradation of beta-catenin by the destruction complex (R-MMU-195253) | 79 | 36 | 1.99 + | 18.05 | 5.82E-30 | 1.68E-28 |
| Downstream signaling events of B Cell Receptor (BCR) (R-MMU-1168372) | 76 | 34 | 1.92 + | 17.72 | 3.76E-28 | 7.76E-27 |
| APC/C:Cdc20 mediated degradation of mitotic proteins (R-MMU-176409) | 74 | 33 | 1.87 + | 17.67 | 2.54E-27 | 4.75E-26 |
| Activation of APC/C and APC/C:Cdc20 mediated degradation of mitotic proteins (R-MMU-176814) | 75 | 33 | 1.89 + | 17.43 | 3.57E-27 | 6.54E-26 |
| Attenuation phase (R-MMU-3371568) | 14 | 6 | 0.35 + | 16.98 | 6.26E-06 | 5.24E-05 |
| PCP/CE pathway (R-MMU-4086400) | 87 | 37 | 2.2 + | 16.85 | 6.63E-30 | 1.88E-28 |
| Regulation of localization of FOXO transcription factors (R-MMU-9614399) | 12 | 5 | 0.3 + | 16.51 | 4.30E-05 | 3.27E-04 |
| Hedgehog 'on' state (R-MMU-5632684) | 82 | 34 | 2.07 + | 16.43 | 2.78E-27 | 5.15E-26 |
| Transcriptional regulation by RUNX3 (R-MMU-8878159) | 81 | 33 | 2.04 + | 16.14 | 2.56E-26 | 4.35E-25 |
| Translation (R-MMU-72766) | 225 | 91 | 5.68 + | 16.02 | 2.43E-71 | 5.72E-69 |
| Regulation of mitotic cell cycle (R-MMU-453276) | 82 | 33 | 2.07 + | 15.94 | 3.51E-26 | 5.90E-25 |
| APC/C-mediated degradation of cell cycle proteins (R-MMU-174143) | 82 | 33 | 2.07 + | 15.94 | 3.51E-26 | 5.84E-25 |
| N-glycan trimming in the ER and Calnexin/Calreticulin cycle (R-MMU-532668) | 15 | 6 | 0.38 + | 15.85 | 8.59E-06 | 7.01E-05 |
| RHO GTPases activate PAKs (R-MMU-5627123) | 20 | 8 | 0.5 + | 15.85 | 2.59E-07 | 2.37E-06 |
| DNA Replication Pre-Initiation (R-MMU-69002) | 83 | 33 | 2.1 + | 15.75 | 4.79E-26 | 7.90E-25 |
| CLEC7A (Dectin-1) signaling (R-MMU-5607764) | 87 | 34 | 2.2 + | 15.48 | 1.33E-26 | 2.37E-25 |
| Activation of BH3-only proteins (R-MMU-114452) | 18 | 7 | 0.45 + | 15.41 | 1.73E-06 | 1.51E-05 |
| Apoptosis induced DNA fragmentation (R-MMU-140342) | 13 | 5 | 0.33 + | 15.24 | 5.84E-05 | 4.37E-04 |
| ERKs are inactivated (R-MMU-202670) | 13 | 5 | 0.33 + | 15.24 | 5.84E-05 | 4.35E-04 |
| Switching of origins to a post-replicative state (R-MMU-69052) | 89 | 34 | 2.25 + | 15.13 | 2.44E-26 | 4.20E-25 |
| ABC-family proteins mediated transport (R-MMU-382556) | 100 | 38 | 2.52 + | 15.05 | 3.11E-29 | 7.43E-28 |
| HSP90 chaperone cycle for steroid hormone receptors (SHR) (R-MMU-3371497) | 53 | 20 | 1.34 + | 14.95 | 6.13E-16 | 7.22E-15 |
| Glycogen synthesis (R-MMU-3322077) | 8 | 3 | 0.2 + | 14.86 | 2.11E-03 | 1.28E-02 |
| RHO GTPases activate CIT (R-MMU-5625900) | 8 | 3 | 0.2 + | 14.86 | 2.11E-03 | 1.27E-02 |
| Recycling of eIF2:GDP (R-MMU-72731) | 8 | 3 | 0.2 + | 14.86 | 2.11E-03 | 1.27E-02 |
| Vitamin C (ascorbate) metabolism (R-MMU-196836) | 8 | 3 | 0.2 + | 14.86 | 2.11E-03 | 1.26E-02 |
| UCH proteinases (R-MMU-5689603) | 95 | 35 | 2.4 + | 14.6 | 1.20E-26 | 2.16E-25 |
| Hedgehog 'off' state (R-MMU-5610787) | 106 | 39 | 2.68 + | 14.58 | 1.52E-29 | 3.98E-28 |
| Interleukin-1 signaling (R-MMU-9020702) | 96 | 35 | 2.42 + | 14.44 | 1.60E-26 | 2.81E-25 |
| RHO GTPases activate KTN1 (R-MMU-5625970) | 11 | 4 | 0.28 + | 14.41 | 4.00E-04 | 2.72E-03 |

| | | | | | | |
|---|---|---|---|---|---|---|
| Antigen processing-Cross presentation (R-MMU-1236975) | 97 | 34 | 2.45 + | 13.89 | 2.44E-25 | 3.98E-24 |
| Major pathway of rRNA processing in the nucleolus and cytosol (R-MMU-6791226) | 175 | 61 | 4.42 + | 13.81 | 1.29E-44 | 1.25E-42 |
| rRNA processing in the nucleus and cytosol (R-MMU-8868773) | 175 | 61 | 4.42 + | 13.81 | 1.29E-44 | 1.18E-42 |
| rRNA processing (R-MMU-72312) | 175 | 61 | 4.42 + | 13.81 | 1.29E-44 | 1.12E-42 |
| Downstream TCR signaling (R-MMU-202424) | 96 | 33 | 2.42 + | 13.62 | 2.08E-24 | 3.17E-23 |
| TNFR2 non-canonical NF-kB pathway (R-MMU-5668541) | 99 | 34 | 2.5 + | 13.61 | 4.21E-25 | 6.68E-24 |
| RHO GTPases activate IQGAPs (R-MMU-5626467) | 27 | 9 | 0.68 + | 13.21 | 1.62E-07 | 1.51E-06 |
| G1/S Transition (R-MMU-69206) | 106 | 35 | 2.68 + | 13.08 | 2.48E-25 | 4.00E-24 |
| C-type lectin receptors (CLRs) (R-MMU-5621481) | 107 | 35 | 2.7 + | 12.96 | 3.21E-25 | 5.14E-24 |
| COPI-independent Golgi-to-ER retrograde traffic (R-MMU-6811436) | 49 | 16 | 1.24 + | 12.94 | 3.14E-12 | 3.50E-11 |
| Beta-catenin independent WNT signaling (R-MMU-3858494) | 126 | 41 | 3.18 + | 12.89 | 2.81E-29 | 6.92E-28 |
| Interleukin-1 family signaling (R-MMU-446652) | 110 | 35 | 2.78 + | 12.61 | 6.92E-25 | 1.08E-23 |
| Cooperation of PDCL (PhLP1) and TRiC/CCT in G-protein beta folding (R-MMU-6814122) | 42 | 13 | 1.06 + | 12.26 | 5.99E-10 | 6.46E-09 |
| RHO GTPases Activate WASPs and WAVEs (R-MMU-5663213) | 36 | 11 | 0.91 + | 12.11 | 1.43E-08 | 1.44E-07 |
| BBSome-mediated cargo-targeting to cilium (R-MMU-5620922) | 23 | 7 | 0.58 + | 12.06 | 6.57E-06 | 5.47E-05 |
| Protein folding (R-MMU-391251) | 43 | 13 | 1.09 + | 11.98 | 7.63E-10 | 8.12E-09 |
| Chaperonin-mediated protein folding (R-MMU-390466) | 43 | 13 | 1.09 + | 11.98 | 7.63E-10 | 8.07E-09 |
| PTEN Regulation (R-MMU-6807070) | 113 | 34 | 2.85 + | 11.92 | 1.49E-23 | 2.13E-22 |
| Formation of annular gap junctions (R-MMU-196025) | 10 | 3 | 0.25 + | 11.89 | 3.53E-03 | 2.02E-02 |
| Golgi Cisternae Pericentriolar Stack Reorganization (R-MMU-162658) | 10 | 3 | 0.25 + | 11.89 | 3.53E-03 | 2.01E-02 |
| G2/M Checkpoints (R-MMU-69481) | 147 | 44 | 3.71 + | 11.86 | 4.47E-30 | 1.37E-28 |
| Signaling by Hedgehog (R-MMU-5358351) | 138 | 41 | 3.48 + | 11.77 | 5.45E-28 | 1.11E-26 |
| TCR signaling (R-MMU-202403) | 115 | 34 | 2.9 + | 11.71 | 2.39E-23 | 3.37E-22 |
| Protein methylation (R-MMU-8876725) | 17 | 5 | 0.43 + | 11.65 | 1.65E-04 | 1.19E-03 |
| HSF1-dependent transactivation (R-MMU-3371571) | 24 | 7 | 0.61 + | 11.56 | 8.31E-06 | 6.85E-05 |
| Synthesis of DNA (R-MMU-69239) | 117 | 34 | 2.95 + | 11.51 | 3.81E-23 | 5.32E-22 |
| Recycling pathway of L1 (R-MMU-437239) | 35 | 10 | 0.88 + | 11.32 | 1.11E-07 | 1.07E-06 |
| Aggrephagy (R-MMU-9646399) | 35 | 10 | 0.88 + | 11.32 | 1.11E-07 | 1.06E-06 |
| EPHB-mediated forward signaling (R-MMU-3928662) | 39 | 11 | 0.98 + | 11.17 | 2.86E-08 | 2.86E-07 |
| G2/M Transition (R-MMU-69275) | 182 | 51 | 4.59 + | 11.1 | 1.49E-33 | 8.80E-32 |
| Cell-extracellular matrix interactions (R-MMU-446353) | 18 | 5 | 0.45 + | 11 | 2.07E-04 | 1.47E-03 |
| Mitotic G2-G2/M phases (R-MMU-453274) | 184 | 51 | 4.64 + | 10.98 | 2.33E-33 | 1.33E-31 |
| Unfolded Protein Response (UPR) (R-MMU-381119) | 11 | 3 | 0.28 + | 10.8 | 4.41E-03 | 2.40E-02 |
| Josephin domain DUBs (R-MMU-5689877) | 11 | 3 | 0.28 + | 10.8 | 4.41E-03 | 2.39E-02 |
| Aspartate and asparagine metabolism (R-MMU-8963693) | 11 | 3 | 0.28 + | 10.8 | 4.41E-03 | 2.39E-02 |
| Gap junction degradation (R-MMU-190873) | 11 | 3 | 0.28 + | 10.8 | 4.41E-03 | 2.38E-02 |
| DNA Replication (R-MMU-69306) | 125 | 34 | 3.16 + | 10.78 | 2.29E-22 | 3.12E-21 |
| Mitotic G1 phase and G1/S transition (R-MMU-453279) | 130 | 35 | 3.28 + | 10.67 | 7.25E-23 | 1.01E-21 |
| Microtubule-dependent trafficking of connexons from Golgi to the plasma membrane (R-MMU-190840) | 15 | 4 | 0.38 + | 10.56 | 1.05E-03 | 6.71E-03 |
| Synthesis of active ubiquitin: roles of E1 and E2 enzymes (R-MMU-8866652) | 30 | 8 | 0.76 + | 10.56 | 3.28E-06 | 2.81E-05 |
| ERK/MAPK targets (R-MMU-198753) | 19 | 5 | 0.48 + | 10.43 | 2.56E-04 | 1.80E-03 |
| Separation of Sister Chromatids (R-MMU-2467813) | 183 | 48 | 4.62 + | 10.39 | 1.59E-30 | 5.45E-29 |
| Frs2-mediated activation (R-MMU-170968) | 12 | 3 | 0.3 + | 9.9 | 5.42E-03 | 2.86E-02 |
| CD28 dependent Vav1 pathway (R-MMU-389359) | 12 | 3 | 0.3 + | 9.9 | 5.42E-03 | 2.85E-02 |
| Spry regulation of FGF signaling (R-MMU-1295596) | 16 | 4 | 0.4 + | 9.9 | 1.29E-03 | 8.04E-03 |
| Mitotic Anaphase (R-MMU-68882) | 192 | 48 | 4.85 + | 9.9 | 1.00E-29 | 2.76E-28 |
| Transport of connexons to the plasma membrane (R-MMU-190872) | 16 | 4 | 0.4 + | 9.9 | 1.29E-03 | 8.01E-03 |
| COPI-mediated anterograde transport (R-MMU-6807878) | 96 | 24 | 2.42 + | 9.9 | 1.82E-15 | 2.10E-14 |
| G beta:gamma signalling through CDC42 (R-MMU-8964616) | 20 | 5 | 0.5 + | 9.9 | 3.14E-04 | 2.17E-03 |
| Mitotic Metaphase and Anaphase (R-MMU-2555396) | 193 | 48 | 4.87 + | 9.85 | 1.22E-29 | 3.31E-28 |
| FCERI mediated NF-kB activation (R-MMU-2871837) | 133 | 33 | 3.36 + | 9.83 | 1.08E-20 | 1.39E-19 |
| Gluconeogenesis (R-MMU-70263) | 34 | 8 | 0.86 + | 9.32 | 7.25E-06 | 6.01E-05 |
| Insertion of tail-anchored proteins into the endoplasmic reticulum membrane (R-MMU-9609523) | 17 | 4 | 0.43 + | 9.32 | 1.56E-03 | 9.63E-03 |
| Smooth Muscle Contraction (R-MMU-445355) | 30 | 7 | 0.76 + | 9.24 | 2.86E-05 | 2.25E-04 |
| S Phase (R-MMU-69242) | 146 | 34 | 3.69 + | 9.23 | 1.55E-20 | 1.96E-19 |
| Golgi-to-ER retrograde transport (R-MMU-8856688) | 130 | 30 | 3.28 + | 9.14 | 3.57E-18 | 4.33E-17 |
| Nuclear Events (kinase and transcription factor activation) (R-MMU-198725) | 22 | 5 | 0.56 + | 9 | 4.58E-04 | 3.11E-03 |
| ADP signalling through P2Y purinoceptor 12 (R-MMU-392170) | 22 | 5 | 0.56 + | 9 | 4.58E-04 | 3.09E-03 |
| FOXO-mediated transcription (R-MMU-9614085) | 22 | 5 | 0.56 + | 9 | 4.58E-04 | 3.08E-03 |
| Detoxification of Reactive Oxygen Species (R-MMU-3299685) | 31 | 7 | 0.78 + | 8.95 | 3.44E-05 | 2.66E-04 |
| Ub-specific processing proteases (R-MMU-5689880) | 182 | 41 | 4.59 + | 8.92 | 4.70E-24 | 6.99E-23 |
| RHO GTPases activate PKNs (R-MMU-5625740) | 58 | 13 | 1.46 + | 8.88 | 1.67E-08 | 1.68E-07 |
| Glycogen metabolism (R-MMU-8982491) | 18 | 4 | 0.45 + | 8.8 | 1.87E-03 | 1.14E-02 |
| Transcriptional regulation by RUNX1 (R-MMU-8878171) | 169 | 37 | 4.27 + | 8.67 | 1.88E-21 | 2.50E-20 |
| TCF dependent signaling in response to WNT (R-MMU-201681) | 192 | 42 | 4.85 + | 8.67 | 3.47E-24 | 5.25E-23 |
| Rap1 signalling (R-MMU-392517) | 14 | 3 | 0.35 + | 8.49 | 7.80E-03 | 3.95E-02 |
| Prolonged ERK activation events (R-MMU-169893) | 14 | 3 | 0.35 + | 8.49 | 7.80E-03 | 3.94E-02 |
| Signaling by the B Cell Receptor (BCR) (R-MMU-983705) | 161 | 34 | 4.06 + | 8.37 | 2.19E-19 | 2.74E-18 |
| Prostacyclin signalling through prostacyclin receptor (R-MMU-392851) | 19 | 4 | 0.48 + | 8.34 | 2.22E-03 | 1.32E-02 |
| Intrinsic Pathway for Apoptosis (R-MMU-109606) | 38 | 8 | 0.96 + | 8.34 | 1.47E-05 | 1.17E-04 |

| | | | | | | | |
|---|---|---|---|---|---|---|---|
| Fc epsilon receptor (FCERI) signaling (R-MMU-2454202) | 178 | 37 | 4.49 | + | 8.24 | 8.69E-21 | 1.14E-19 |
| Cell Cycle Checkpoints (R-MMU-69620) | 268 | 55 | 6.76 | + | 8.13 | 3.63E-30 | 1.17E-28 |
| Selective autophagy (R-MMU-9663891) | 59 | 12 | 1.49 | + | 8.06 | 1.51E-07 | 1.41E-06 |
| Pentose phosphate pathway (R-MMU-71336) | 15 | 3 | 0.38 | + | 7.92 | 9.20E-03 | 4.57E-02 |
| Sema4D in semaphorin signaling (R-MMU-400685) | 15 | 3 | 0.38 | + | 7.92 | 9.20E-03 | 4.55E-02 |
| Antiviral mechanism by IFN-stimulated genes (R-MMU-1169410) | 35 | 7 | 0.88 | + | 7.92 | 6.74E-05 | 5.01E-04 |
| Negative regulation of MAPK pathway (R-MMU-5675221) | 40 | 8 | 1.01 | + | 7.92 | 2.04E-05 | 1.61E-04 |
| MAP2K and MAPK activation (R-MMU-5674135) | 40 | 8 | 1.01 | + | 7.92 | 2.04E-05 | 1.61E-04 |
| Metabolism of RNA (R-MMU-8953854) | 566 | 112 | 14.29 | + | 7.84 | 2.72E-60 | 3.73E-58 |
| Cellular responses to stress (R-MMU-2262752) | 411 | 81 | 10.37 | + | 7.81 | 3.80E-43 | 3.13E-41 |
| L1CAM interactions (R-MMU-373760) | 66 | 13 | 1.67 | + | 7.8 | 6.35E-08 | 6.16E-07 |
| Cellular responses to external stimuli (R-MMU-8953897) | 413 | 81 | 10.42 | + | 7.77 | 5.21E-43 | 4.09E-41 |
| MAPK family signaling cascades (R-MMU-5683057) | 271 | 53 | 6.84 | + | 7.75 | 3.27E-28 | 6.82E-27 |
| RAF/MAP kinase cascade (R-MMU-5673001) | 241 | 47 | 6.08 | + | 7.73 | 4.49E-25 | 7.06E-24 |
| ISG15 antiviral mechanism (R-MMU-1169408) | 31 | 6 | 0.78 | + | 7.67 | 2.63E-04 | 1.84E-03 |
| Signal transduction by L1 (R-MMU-445144) | 21 | 4 | 0.53 | + | 7.55 | 3.05E-03 | 1.78E-02 |
| MAPK1/MAPK3 signaling (R-MMU-5684996) | 247 | 47 | 6.23 | + | 7.54 | 1.13E-24 | 1.74E-23 |
| COPI-dependent Golgi-to-ER retrograde traffic (R-MMU-6811434) | 95 | 18 | 2.4 | + | 7.51 | 3.26E-10 | 3.54E-09 |
| Protein ubiquitination (R-MMU-8852135) | 64 | 12 | 1.62 | + | 7.43 | 3.26E-07 | 2.97E-06 |
| G-protein beta:gamma signalling (R-MMU-397795) | 32 | 6 | 0.81 | + | 7.43 | 3.06E-04 | 2.12E-03 |
| Signaling by WNT (R-MMU-195721) | 273 | 51 | 6.89 | + | 7.4 | 2.26E-26 | 3.93E-25 |
| Neddylation (R-MMU-8951664) | 225 | 42 | 5.68 | + | 7.4 | 6.99E-22 | 9.37E-21 |
| Loss of proteins required for interphase microtubule organization from the centrosome (R-MMU-380284) | 70 | 13 | 1.77 | + | 7.36 | 1.16E-07 | 1.11E-06 |
| Loss of Nlp from mitotic centrosomes (R-MMU-380259) | 70 | 13 | 1.77 | + | 7.36 | 1.16E-07 | 1.10E-06 |
| TP53 Regulates Metabolic Genes (R-MMU-5628897) | 49 | 9 | 1.24 | + | 7.28 | 1.13E-05 | 9.17E-05 |
| FLT3 Signaling (R-MMU-9607240) | 256 | 47 | 6.46 | + | 7.27 | 4.29E-24 | 6.43E-23 |
| Deubiquitination (R-MMU-5688426) | 251 | 46 | 6.34 | + | 7.26 | 1.42E-23 | 2.05E-22 |
| RAF-independent MAPK1/3 activation (R-MMU-112409) | 22 | 4 | 0.56 | + | 7.2 | 3.53E-03 | 2.01E-02 |
| ER to Golgi Anterograde Transport (R-MMU-199977) | 149 | 27 | 3.76 | + | 7.18 | 3.11E-14 | 3.53E-13 |
| Apoptotic execution phase (R-MMU-75153) | 50 | 9 | 1.26 | + | 7.13 | 1.31E-05 | 1.05E-04 |
| G-protein activation (R-MMU-202040) | 28 | 5 | 0.71 | + | 7.07 | 1.19E-03 | 7.56E-03 |
| MAPK targets/ Nuclear events mediated by MAP kinases (R-MMU-450282) | 28 | 5 | 0.71 | + | 7.07 | 1.19E-03 | 7.53E-03 |
| Pyruvate metabolism (R-MMU-70268) | 28 | 5 | 0.71 | + | 7.07 | 1.19E-03 | 7.50E-03 |
| AURKA Activation by TPX2 (R-MMU-8854518) | 73 | 13 | 1.84 | + | 7.06 | 1.79E-07 | 1.66E-06 |
| Adrenaline,noradrenaline inhibits insulin secretion (R-MMU-400042) | 23 | 4 | 0.58 | + | 6.89 | 4.07E-03 | 2.24E-02 |
| EPH-Ephrin signaling (R-MMU-2682334) | 70 | 12 | 1.77 | + | 6.79 | 7.62E-07 | 6.76E-06 |
| Gap junction trafficking (R-MMU-190828) | 41 | 7 | 1.03 | + | 6.76 | 1.62E-04 | 1.17E-03 |
| Cellular response to heat stress (R-MMU-3371556) | 88 | 15 | 2.22 | + | 6.75 | 3.42E-08 | 3.39E-07 |
| Intra-Golgi and retrograde Golgi-to-ER traffic (R-MMU-6811442) | 188 | 32 | 4.75 | + | 6.74 | 6.06E-16 | 7.19E-15 |
| M Phase (R-MMU-68886) | 354 | 60 | 8.94 | + | 6.71 | 7.02E-29 | 1.56E-27 |
| RAF activation (R-MMU-5673000) | 30 | 5 | 0.76 | + | 6.6 | 1.57E-03 | 9.64E-03 |
| Recruitment of NuMA to mitotic centrosomes (R-MMU-380320) | 90 | 15 | 2.27 | + | 6.6 | 4.45E-08 | 4.37E-07 |
| Metabolism of amino acids and derivatives (R-MMU-71291) | 252 | 42 | 6.36 | + | 6.6 | 3.01E-20 | 3.79E-19 |
| RHO GTPase Effectors (R-MMU-195258) | 284 | 47 | 7.17 | + | 6.56 | 2.03E-22 | 2.79E-21 |
| Signaling by Interleukins (R-MMU-449147) | 260 | 43 | 6.56 | + | 6.55 | 1.37E-20 | 1.75E-19 |
| PIP3 activates AKT signaling (R-MMU-1257604) | 238 | 39 | 6.01 | + | 6.49 | 1.17E-18 | 1.43E-17 |
| Apoptosis (R-MMU-109581) | 104 | 17 | 2.63 | + | 6.48 | 7.31E-09 | 7.44E-08 |
| Gap junction trafficking and regulation (R-MMU-157858) | 43 | 7 | 1.09 | + | 6.45 | 2.11E-04 | 1.48E-03 |
| Centrosome maturation (R-MMU-380287) | 80 | 13 | 2.02 | + | 6.44 | 4.56E-07 | 4.09E-06 |
| Recruitment of mitotic centrosome proteins and complexes (R-MMU-380270) | 80 | 13 | 2.02 | + | 6.44 | 4.56E-07 | 4.07E-06 |
| Programmed Cell Death (R-MMU-5357801) | 111 | 18 | 2.8 | + | 6.42 | 2.96E-09 | 3.07E-08 |
| Regulation of PLK1 Activity at G2/M Transition (R-MMU-2565942) | 87 | 14 | 2.2 | + | 6.38 | 1.84E-07 | 1.70E-06 |
| Cargo trafficking to the periciliary membrane (R-MMU-5620920) | 50 | 8 | 1.26 | + | 6.34 | 8.33E-05 | 6.16E-04 |
| G beta:gamma signalling through PI3Kgamma (R-MMU-392451) | 25 | 4 | 0.63 | + | 6.34 | 5.30E-03 | 2.81E-02 |
| DNA Damage Recognition in GG-NER (R-MMU-5696394) | 38 | 6 | 0.96 | + | 6.26 | 6.91E-04 | 4.52E-03 |
| RHO GTPases Activate Formins (R-MMU-5663220) | 133 | 21 | 3.36 | + | 6.26 | 2.21E-10 | 2.43E-09 |
| Formation of the cornified envelope (R-MMU-6809371) | 121 | 19 | 3.05 | + | 6.22 | 1.74E-09 | 1.81E-08 |
| Vasopressin regulates renal water homeostasis via Aquaporins (R-MMU-432040) | 32 | 5 | 0.81 | + | 6.19 | 2.02E-03 | 1.23E-02 |
| Signal amplification (R-MMU-392518) | 32 | 5 | 0.81 | + | 6.19 | 2.02E-03 | 1.22E-02 |
| Intracellular signaling by second messengers (R-MMU-9006925) | 264 | 41 | 6.66 | + | 6.15 | 8.31E-19 | 1.02E-17 |
| Signaling by NTRK1 (TRKA) (R-MMU-187037) | 58 | 9 | 1.46 | + | 6.15 | 3.74E-05 | 2.87E-04 |
| Antigen processing: Ubiquitination & Proteasome degradation (R-MMU-983168) | 297 | 46 | 7.5 | + | 6.14 | 6.33E-21 | 8.36E-20 |
| Carboxyterminal post-translational modifications of tubulin (R-MMU-8955332) | 26 | 4 | 0.66 | + | 6.09 | 5.99E-03 | 3.14E-02 |
| Transport to the Golgi and subsequent modification (R-MMU-948021) | 179 | 27 | 4.52 | + | 5.98 | 1.51E-12 | 1.69E-11 |
| MHC class II antigen presentation (R-MMU-2132295) | 133 | 20 | 3.36 | + | 5.96 | 1.27E-09 | 1.33E-08 |
| Deadenylation of mRNA (R-MMU-429947) | 27 | 4 | 0.68 | + | 5.87 | 6.75E-03 | 3.47E-02 |
| VEGFR2 mediated vascular permeability (R-MMU-5218920) | 27 | 4 | 0.68 | + | 5.87 | 6.75E-03 | 3.46E-02 |

| | | | | | | | |
|---|---|---|---|---|---|---|---|
| Lysosome Vesicle Biogenesis (R-MMU-432720) | 34 | 5 | 0.86 | + | 5.83 | 2.56E-03 | 1.50E-02 |
| G2/M DNA damage checkpoint (R-MMU-69473) | 75 | 11 | 1.89 | + | 5.81 | 8.42E-06 | 6.91E-05 |
| Class I MHC mediated antigen processing & presentation (R-MMU-983169) | 361 | 52 | 9.11 | + | 5.71 | 3.00E-22 | 4.05E-21 |
| Asparagine N-linked glycosylation (R-MMU-446203) | 265 | 38 | 6.69 | + | 5.68 | 1.71E-16 | 2.05E-15 |
| Signaling by NTRKs (R-MMU-166520) | 71 | 10 | 1.79 | + | 5.58 | 2.96E-05 | 2.31E-04 |
| Opioid Signalling (R-MMU-111885) | 71 | 10 | 1.79 | + | 5.58 | 2.96E-05 | 2.30E-04 |
| EML4 and NUDC in mitotic spindle formation (R-MMU-9648025) | 108 | 15 | 2.73 | + | 5.5 | 3.73E-07 | 3.38E-06 |
| Negative regulation of FGFR3 signaling (R-MMU-5654732) | 29 | 4 | 0.73 | + | 5.46 | 8.45E-03 | 4.23E-02 |
| Glucose metabolism (R-MMU-70326) | 80 | 11 | 2.02 | + | 5.45 | 1.46E-05 | 1.17E-04 |
| E3 ubiquitin ligases ubiquitinate target proteins (R-MMU-8866654) | 44 | 6 | 1.11 | + | 5.4 | 1.37E-03 | 8.52E-03 |
| Neutrophil degranulation (R-MMU-6798695) | 548 | 74 | 13.83 | + | 5.35 | 8.55E-30 | 2.39E-28 |
| Pyruvate metabolism and Citric Acid (TCA) cycle (R-MMU-71406) | 52 | 7 | 1.31 | + | 5.33 | 5.95E-04 | 3.92E-03 |
| Activation of kainate receptors upon glutamate binding (R-MMU-451326) | 30 | 4 | 0.76 | + | 5.28 | 9.39E-03 | 4.64E-02 |
| Anchoring of the basal body to the plasma membrane (R-MMU-5620912) | 98 | 13 | 2.47 | + | 5.26 | 3.53E-06 | 3.01E-05 |
| Regulation of HSF1-mediated heat shock response (R-MMU-3371453) | 68 | 9 | 1.72 | + | 5.24 | 1.14E-04 | 8.29E-04 |
| Signaling by Rho GTPases (R-MMU-194315) | 410 | 53 | 10.35 | + | 5.12 | 9.72E-21 | 1.26E-19 |
| Aquaporin-mediated transport (R-MMU-445717) | 39 | 5 | 0.98 | + | 5.08 | 4.37E-03 | 2.39E-02 |
| PLC beta mediated events (R-MMU-112043) | 39 | 5 | 0.98 | + | 5.08 | 4.37E-03 | 2.39E-02 |
| Metabolism of proteins (R-MMU-392499) | 1644 | 210 | 41.5 | + | 5.06 | 2.38E-86 | 3.93E-83 |
| Cell Cycle, Mitotic (R-MMU-69278) | 480 | 61 | 12.12 | + | 5.03 | 2.11E-23 | 3.00E-22 |
| Cilium Assembly (R-MMU-5617833) | 198 | 25 | 5 | + | 5 | 3.11E-10 | 3.39E-09 |
| Resolution of Sister Chromatid Cohesion (R-MMU-2500257) | 120 | 15 | 3.03 | + | 4.95 | 1.25E-06 | 1.10E-05 |
| G-protein mediated events (R-MMU-112040) | 40 | 5 | 1.01 | + | 4.95 | 4.82E-03 | 2.58E-02 |
| MAP kinase activation (R-MMU-450294) | 57 | 7 | 1.44 | + | 4.87 | 9.75E-04 | 6.28E-03 |
| Interleukin-17 signaling (R-MMU-448424) | 57 | 7 | 1.44 | + | 4.87 | 9.75E-04 | 6.25E-03 |
| Cell Cycle (R-MMU-1640170) | 552 | 66 | 13.93 | + | 4.74 | 6.14E-24 | 9.04E-23 |
| Amplification of signal from unattached kinetochores via a MAD2 inhibitory signal (R-MMU-141444) | 93 | 11 | 2.35 | + | 4.69 | 5.16E-05 | 3.90E-04 |
| Amplification of signal from the kinetochores (R-MMU-141424) | 93 | 11 | 2.35 | + | 4.69 | 5.16E-05 | 3.88E-04 |
| Cell junction organization (R-MMU-446728) | 68 | 8 | 1.72 | + | 4.66 | 5.57E-04 | 3.72E-03 |
| COPII-mediated vesicle transport (R-MMU-204005) | 68 | 8 | 1.72 | + | 4.66 | 5.57E-04 | 3.70E-03 |
| Glycolysis (R-MMU-70171) | 60 | 7 | 1.51 | + | 4.62 | 1.28E-03 | 8.03E-03 |
| Formation of Incision Complex in GG-NER (R-MMU-5696395) | 43 | 5 | 1.09 | + | 4.61 | 6.36E-03 | 3.30E-02 |
| Extra-nuclear estrogen signaling (R-MMU-9009391) | 69 | 8 | 1.74 | + | 4.59 | 6.09E-04 | 4.00E-03 |
| Intraflagellar transport (R-MMU-5620924) | 53 | 6 | 1.34 | + | 4.48 | 3.24E-03 | 1.87E-02 |
| Integration of energy metabolism (R-MMU-163685) | 80 | 9 | 2.02 | + | 4.46 | 3.45E-04 | 2.37E-03 |
| Organelle biogenesis and maintenance (R-MMU-1852241) | 224 | 25 | 5.65 | + | 4.42 | 3.14E-09 | 3.23E-08 |
| Innate Immune System (R-MMU-168249) | 1032 | 115 | 26.05 | + | 4.41 | 2.87E-39 | 2.06E-37 |
| Golgi Associated Vesicle Biogenesis (R-MMU-432722) | 55 | 6 | 1.39 | + | 4.32 | 3.83E-03 | 2.12E-02 |
| Recruitment and ATM-mediated phosphorylation of repair and signaling proteins at DNA double strand breaks (R-MMU-5693565) | 56 | 6 | 1.41 | + | 4.24 | 4.15E-03 | 2.28E-02 |
| Cytokine Signaling in Immune system (R-MMU-1280215) | 544 | 58 | 13.73 | + | 4.22 | 5.84E-19 | 7.24E-18 |
| DNA Double Strand Break Response (R-MMU-5693606) | 57 | 6 | 1.44 | + | 4.17 | 4.49E-03 | 2.41E-02 |
| Platelet homeostasis (R-MMU-418346) | 76 | 8 | 1.92 | + | 4.17 | 1.09E-03 | 6.91E-03 |
| DNA Damage/Telomere Stress Induced Senescence (R-MMU-2559586) | 67 | 7 | 1.69 | + | 4.14 | 2.29E-03 | 1.35E-02 |
| Mitotic Prometaphase (R-MMU-68877) | 194 | 20 | 4.9 | + | 4.08 | 3.80E-07 | 3.42E-06 |
| Macroautophagy (R-MMU-1632852) | 117 | 12 | 2.95 | + | 4.06 | 8.45E-05 | 6.22E-04 |
| Autophagy (R-MMU-9612973) | 117 | 12 | 2.95 | + | 4.06 | 8.45E-05 | 6.20E-04 |
| Mitotic Spindle Checkpoint (R-MMU-69618) | 109 | 11 | 2.75 | + | 4 | 1.89E-04 | 1.35E-03 |
| Adaptive Immune System (R-MMU-1280218) | 780 | 78 | 19.69 | + | 3.96 | 1.06E-23 | 1.55E-22 |
| Cell-Cell communication (R-MMU-1500931) | 90 | 9 | 2.27 | + | 3.96 | 7.57E-04 | 4.93E-03 |
| Toll Like Receptor 3 (TLR3) Cascade (R-MMU-168164) | 70 | 7 | 1.77 | + | 3.96 | 2.87E-03 | 1.68E-02 |
| Regulation of insulin secretion (R-MMU-422356) | 61 | 6 | 1.54 | + | 3.9 | 6.08E-03 | 3.16E-02 |
| Interferon Signaling (R-MMU-913531) | 72 | 7 | 1.82 | + | 3.85 | 3.32E-03 | 1.91E-02 |
| MyD88 cascade initiated on plasma membrane (R-MMU-975871) | 73 | 7 | 1.84 | + | 3.8 | 3.56E-03 | 2.02E-02 |
| Toll Like Receptor 5 (TLR5) Cascade (R-MMU-168176) | 73 | 7 | 1.84 | + | 3.8 | 3.56E-03 | 2.01E-02 |
| Toll Like Receptor 10 (TLR10) Cascade (R-MMU-168142) | 73 | 7 | 1.84 | + | 3.8 | 3.56E-03 | 2.00E-02 |
| Membrane Trafficking (R-MMU-199991) | 564 | 54 | 14.24 | + | 3.79 | 7.09E-16 | 8.29E-15 |
| Regulation of actin dynamics for phagocytic cup formation (R-MMU-2029482) | 126 | 12 | 3.18 | + | 3.77 | 1.61E-04 | 1.17E-03 |
| Toll Like Receptor 2 (TLR2) Cascade (R-MMU-181438) | 74 | 7 | 1.87 | + | 3.75 | 3.82E-03 | 2.14E-02 |
| MyD88:MAL(TIRAP) cascade initiated on plasma membrane (R-MMU-166058) | 74 | 7 | 1.87 | + | 3.75 | 3.82E-03 | 2.13E-02 |
| Toll Like Receptor TLR6:TLR2 Cascade (R-MMU-168188) | 74 | 7 | 1.87 | + | 3.75 | 3.82E-03 | 2.13E-02 |
| Toll Like Receptor TLR1:TLR2 Cascade (R-MMU-168179) | 74 | 7 | 1.87 | + | 3.75 | 3.82E-03 | 2.12E-02 |
| Axon guidance (R-MMU-422475) | 277 | 26 | 6.99 | + | 3.72 | 3.99E-08 | 3.94E-07 |
| Factors involved in megakaryocyte development and platelet production (R-MMU-983231) | 119 | 11 | 3 | + | 3.66 | 3.79E-04 | 2.59E-03 |
| ESR-mediated signaling (R-MMU-8939211) | 163 | 15 | 4.11 | + | 3.65 | 3.67E-05 | 2.83E-04 |
| Clathrin-mediated endocytosis (R-MMU-8856828) | 142 | 13 | 3.58 | + | 3.63 | 1.25E-04 | 9.11E-04 |
| Transport of small molecules (R-MMU-382551) | 674 | 61 | 17.01 | + | 3.59 | 9.41E-17 | 1.13E-15 |
| Keratinization (R-MMU-6805567) | 212 | 19 | 5.35 | + | 3.55 | 4.98E-06 | 4.21E-05 |

| | | | | | | | |
|---|---|---|---|---|---|---|---|
| VEGFA-VEGFR2 Pathway (R-MMU-4420097) | 91 | 8 | 2.3 | + | 3.48 | 3.10E-03 | 1.80E-02 |
| Global Genome Nucleotide Excision Repair (GG-NER) (R-MMU-5696399) | 81 | 7 | 2.04 | + | 3.42 | 6.02E-03 | 3.14E-02 |
| TRAF6 mediated induction of NFkB and MAP kinases upon TLR7/8 or 9 activation (R-MMU-975138) | 83 | 7 | 2.1 | + | 3.34 | 6.80E-03 | 3.47E-02 |
| Post-translational protein modification (R-MMU-597592) | 1283 | 108 | 32.38 | + | 3.33 | 4.36E-27 | 7.91E-26 |
| Protein localization (R-MMU-9609507) | 107 | 9 | 2.7 | + | 3.33 | 2.31E-03 | 1.36E-02 |
| Platelet activation, signaling and aggregation (R-MMU-76002) | 251 | 21 | 6.34 | + | 3.31 | 4.35E-06 | 3.70E-05 |
| Developmental Biology (R-MMU-1266738) | 538 | 45 | 13.58 | + | 3.31 | 1.49E-11 | 1.65E-10 |
| Toll Like Receptor 7/8 (TLR7/8) Cascade (R-MMU-168181) | 84 | 7 | 2.12 | + | 3.3 | 7.21E-03 | 3.67E-02 |
| MyD88 dependent cascade initiated on endosome (R-MMU-975155) | 84 | 7 | 2.12 | + | 3.3 | 7.21E-03 | 3.66E-02 |
| Metabolism of carbohydrates (R-MMU-71387) | 266 | 22 | 6.71 | + | 3.28 | 3.07E-06 | 2.68E-05 |
| Immune System (R-MMU-168256) | 1854 | 153 | 46.8 | + | 3.27 | 2.43E-38 | 1.67E-36 |
| Vesicle-mediated transport (R-MMU-5653656) | 656 | 54 | 16.56 | + | 3.26 | 2.19E-13 | 2.47E-12 |
| Platelet degranulation (R-MMU-114608) | 122 | 10 | 3.08 | + | 3.25 | 1.61E-03 | 9.88E-03 |
| Fcgamma receptor (FCGR) dependent phagocytosis (R-MMU-2029480) | 147 | 12 | 3.71 | + | 3.23 | 5.93E-04 | 3.92E-03 |
| Signaling by VEGF (R-MMU-194138) | 99 | 8 | 2.5 | + | 3.2 | 4.98E-03 | 2.66E-02 |
| Metabolism of nucleotides (R-MMU-15869) | 100 | 8 | 2.52 | + | 3.17 | 5.26E-03 | 2.80E-02 |
| Toll Like Receptor 9 (TLR9) Cascade (R-MMU-168138) | 88 | 7 | 2.22 | + | 3.15 | 9.06E-03 | 4.51E-02 |
| Response to elevated platelet cytosolic Ca2+ (R-MMU-76005) | 127 | 10 | 3.21 | + | 3.12 | 2.12E-03 | 1.27E-02 |
| MyD88-independent TLR4 cascade (R-MMU-166166) | 90 | 7 | 2.27 | + | 3.08 | 1.01E-02 | 4.97E-02 |
| TRIF(TICAM1)-mediated TLR4 signaling (R-MMU-937061) | 90 | 7 | 2.27 | + | 3.08 | 1.01E-02 | 4.96E-02 |
| Mitotic Prophase (R-MMU-68875) | 104 | 8 | 2.63 | + | 3.05 | 6.53E-03 | 3.37E-02 |
| Signaling by Nuclear Receptors (R-MMU-9006931) | 214 | 16 | 5.4 | + | 2.96 | 1.99E-04 | 1.42E-03 |
| Rho GTPase cycle (R-MMU-194840) | 136 | 10 | 3.43 | + | 2.91 | 3.37E-03 | 1.94E-02 |
| Toll Like Receptor 4 (TLR4) Cascade (R-MMU-166016) | 109 | 8 | 2.75 | + | 2.91 | 8.43E-03 | 4.24E-02 |
| Generic Transcription Pathway (R-MMU-212436) | 816 | 54 | 20.6 | + | 2.62 | 7.47E-10 | 8.00E-09 |
| Hemostasis (R-MMU-109582) | 596 | 36 | 15.04 | + | 2.39 | 5.17E-06 | 4.35E-05 |
| Metabolism (R-MMU-1430728) | 1768 | 105 | 44.63 | + | 2.35 | 1.06E-15 | 1.23E-14 |
| Signaling by Receptor Tyrosine Kinases (R-MMU-9006934) | 415 | 24 | 10.48 | + | 2.29 | 2.70E-04 | 1.88E-03 |
| RNA Polymerase II Transcription (R-MMU-73857) | 936 | 54 | 23.63 | + | 2.29 | 4.68E-08 | 4.57E-07 |
| Gene expression (Transcription) (R-MMU-74160) | 1043 | 57 | 26.33 | + | 2.17 | 1.22E-07 | 1.15E-06 |
| Signal Transduction (R-MMU-162582) | 2486 | 128 | 62.75 | + | 2.04 | 1.23E-14 | 1.40E-13 |
| Unclassified (UNCLASSIFIED) | 12982 | 120 | 327.68 | - | 0.37 | 1.13E-69 | 2.32E-67 |
| G alpha (s) signalling events (R-MMU-418555) | 527 | 4 | 13.3 | - | 0.3 | 6.41E-03 | 3.31E-02 |
| Olfactory Signaling Pathway (R-MMU-381753) | 400 | 1 | 10.1 | - | 0.1 | 9.04E-04 | 5.84E-03 |

**S2 A. 18 placental proteins- differential expressed in GDM-Reactome pathway**

| #term ID | term description | observed gene count | background gene count | FDR | Protein name |
|---|---|---|---|---|---|
| HSA-1474244 | Extracellular matrix organization | 6 | 298 | 1.80E-05 | CTSS,ITGB3,LAMB1,LOXL2,NID1,SPARC |
| HSA-392499 | Metabolism of proteins | 9 | 1948 | 0.00071 | CALR,CTSZ,HDGF,HSPA5,ITM2B,LAMB1,MAN2A1,MFGE8, |
| HSA-3000178 | ECM proteoglycans | 3 | 75 | 0.0016 | ITGB3,LAMB1,SPARC |
| HSA-381119 | Unfolded Protein Response (UPR) | 3 | 94 | 0.0016 | CALR,HDGF,HSPA5 |
| HSA-381183 | ATF6 (ATF6-alpha) activates | 2 | 9 | 0.0016 | CALR,HSPA5 |
| HSA-8957275 | Post-translational protein | 3 | 106 | 0.0019 | LAMB1,MFGE8,SERPINC1 |
| HSA-109582 | Hemostasis | 5 | 601 | 0.002 | HSPA5,ITGB3,SERPINC1,SPARC,SRI |
| HSA-114608 | Platelet degranulation | 3 | 125 | 0.0023 | HSPA5,ITGB3,SPARC |
| HSA-381426 | Regulation of Insulin-like Growth | 3 | 123 | 0.0023 | LAMB1,MFGE8,SERPINC1 |
| HSA-1474228 | Degradation of the extracellular | 3 | 139 | 0.0025 | CTSS,LAMB1,NID1 |
| HSA-983170 | Antigen Presentation: Folding, | 2 | 25 | 0.0025 | CALR,HSPA5 |
| HSA-3000157 | Laminin interactions | 2 | 30 | 0.0031 | LAMB1,NID1 |
| HSA-2173782 | Binding and Uptake of Ligands by | 2 | 40 | 0.005 | CALR,SPARC |
| HSA-1566948 | Elastic fibre formation | 2 | 44 | 0.0057 | ITGB3,LOXL2 |
| HSA-597592 | Post-translational protein | 6 | 1366 | 0.0057 | CALR,CTSZ,LAMB1,MAN2A1,MFGE8,SERPINC1 |
| HSA-381070 | IRE1alpha activates chaperones | 2 | 56 | 0.008 | HDGF,HSPA5 |
| HSA-3000171 | Non-integrin membrane-ECM | 2 | 58 | 0.0081 | ITGB3,LAMB1 |
| HSA-2022090 | Assembly of collagen fibrils and | 2 | 60 | 0.0082 | CTSS,LOXL2 |
| HSA-446203 | Asparagine N-linked glycosylation | 3 | 298 | 0.0118 | CALR,CTSZ,MAN2A1 |
| HSA-5653656 | Vesicle-mediated transport | 4 | 649 | 0.0118 | CALR,CTSZ,MAN2A1,SPARC |
| HSA-977225 | Amyloid fiber formation | 2 | 78 | 0.0118 | ITM2B,MFGE8 |
| HSA-1236975 | Antigen processing-Cross presentation | 2 | 96 | 0.0156 | CALR,CTSS |
| HSA-983169 | Class I MHC mediated antigen processing & presentation | 3 | 365 | 0.0172 | CALR,CTSS,HSPA5 |
| HSA-373760 | L1CAM interactions | 2 | 116 | 0.0208 | ITGB3,LAMB1 |
| HSA-948021 | Transport to the Golgi and subsequent modification | 2 | 181 | 0.0468 | CTSZ,MAN2A1 |

**S2 B. 47 placental proteins- differential expressed in PE-Reactome pathway**

| #term ID | term description | observed | background | false | Protein names |
|---|---|---|---|---|---|
| HSA-168256 | Immune System | 22 | 1925 | 2.80E-08 | ACTG1,ANXA1,ANXA2,COL |
| HSA-114608 | Platelet degranulation | 8 | 125 | 1.32E-07 | A2M,CALU,MANF,PROS1,Q |
| HSA-168249 | Innate Immune System | 15 | 1012 | 6.19E-07 | ACTG1,ANXA2,CTSA,CTSD, |
| HSA-76002 | Platelet activation, signaling and aggregation | 9 | 256 | 6.61E-07 | A2M,CALU,COL1A2,MANF, |
| HSA-6798695 | Neutrophil degranulation | 11 | 471 | 6.68E-07 | ANXA2,CTSA,CTSD,DPP7,G |
| HSA-1474244 | Extracellular matrix organization | 9 | 298 | 1.68E-06 | A2M,COL1A2,COL5A2,CTS |
| HSA-1474228 | Degradation of the extracellular matrix | 7 | 139 | 1.95E-06 | A2M,COL1A2,COL5A2,CTS |
| HSA-109582 | Hemostasis | 11 | 601 | 5.05E-06 | A2M,ANXA2,CALU,COL1A2 |
| HSA-8957275 | Post-translational protein phosphorylation | 5 | 106 | 0.0002 | CALU,CP,HSP90B1,QSOX1, |
| HSA-449147 | Signaling by Interleukins | 8 | 439 | 0.00026 | ANXA1,ANXA2,COL1A2,HS |
| HSA-381426 | Regulation of Insulin-like Growth Factor (IGF) transport and | 5 | 123 | 0.00033 | CALU,CP,HSP90B1,QSOX1, |
| HSA-1442490 | Collagen degradation | 4 | 64 | 0.00047 | COL1A2,COL5A2,CTSD,CTS |
| HSA-3000178 | ECM proteoglycans | 4 | 75 | 0.00079 | COL1A2,COL5A2,DCN,TGF |
| HSA-140877 | Formation of Fibrin Clot (Clotting Cascade) | 3 | 39 | 0.0026 | A2M,PRCP,PROS1 |
| HSA-392499 | Metabolism of proteins | 14 | 1948 | 0.0026 | CALU,CCNA2,COPA,CP,CTS |
| HSA-6785807 | Interleukin-4 and Interleukin-13 signaling | 4 | 106 | 0.0026 | ANXA1,COL1A2,HSP90B1,T |
| HSA-1630316 | Glycosaminoglycan metabolism | 4 | 122 | 0.0036 | DCN,GNS,GUSB,HEXB |
| HSA-2206281 | Mucopolysaccharidoses | 2 | 11 | 0.0063 | GNS,GUSB |
| HSA-2022090 | Assembly of collagen fibrils and other multimeric structures | 3 | 60 | 0.0064 | COL1A2,COL5A2,PCOLCE |
| HSA-2160916 | Hyaluronan uptake and degradation | 2 | 12 | 0.0066 | GUSB,HEXB |
| HSA-1650814 | Collagen biosynthesis and modifying enzymes | 3 | 67 | 0.0073 | COL1A2,COL5A2,PCOLCE |
| HSA-1679131 | Trafficking and processing of endosomal TLR | 2 | 13 | 0.0073 | CTSK,HSP90B1 |
| HSA-2022857 | Keratan sulfate degradation | 2 | 13 | 0.0073 | GNS,HEXB |
| HSA-2024101 | CS/DS degradation | 2 | 14 | 0.0074 | DCN,HEXB |
| HSA-2243919 | Crosslinking of collagen fibrils | 2 | 18 | 0.0108 | COL1A2,PCOLCE |
| HSA-3000480 | Scavenging by Class A Receptors | 2 | 18 | 0.0108 | COL1A2,HSP90B1 |
| HSA-597592 | Post-translational protein modification | 10 | 1366 | 0.0128 | CALU,CCNA2,COPA,CP,CTS |
| HSA-140837 | Intrinsic Pathway of Fibrin Clot Formation | 2 | 22 | 0.0136 | A2M,PRCP |
| HSA-3000170 | Syndecan interactions | 2 | 26 | 0.0179 | COL1A2,COL5A2 |
| HSA-8874081 | MET activates PTK2 signaling | 2 | 29 | 0.0213 | COL1A2,COL5A2 |
| HSA-1643685 | Disease | 8 | 1018 | 0.0231 | ACTG1,CP,DCN,GNS,GUSB, |
| HSA-445355 | Smooth Muscle Contraction | 2 | 32 | 0.0242 | ANXA1,ANXA2 |
| HSA-1592389 | Activation of Matrix Metalloproteinases | 2 | 33 | 0.0244 | CTSK,TIMP1 |
| HSA-2132295 | MHC class II antigen presentation | 3 | 119 | 0.0244 | CTSA,CTSD,CTSK |
| HSA-3299685 | Detoxification of Reactive Oxygen Species | 2 | 35 | 0.0251 | GPX3,SOD1 |
| HSA-8950505 | Gene and protein expression by JAK-STAT signaling after | 2 | 38 | 0.0279 | ANXA2,SOD1 |
| HSA-3560782 | Diseases associated with glycosaminoglycan metabolism | 2 | 40 | 0.029 | DCN,HEXB |
| HSA-3781865 | Diseases of glycosylation | 3 | 136 | 0.029 | DCN,HEXB,SPON1 |
| HSA-5653656 | Vesicle-mediated transport | 6 | 649 | 0.029 | ACTG1,COL1A2,COPA,GNS, |
| HSA-1660662 | Glycosphingolipid metabolism | 2 | 44 | 0.0322 | CTSA,HEXB |
| HSA-8948216 | Collagen chain trimerization | 2 | 44 | 0.0322 | COL1A2,COL5A2 |
| HSA-1638091 | Heparan sulfate/heparin (HS-GAG) metabolism | 2 | 55 | 0.0451 | DCN,GUSB |
| HSA-917937 | Iron uptake and transport | 2 | 56 | 0.0458 | CP,TFRC |

**S2 C. 7 placental proteins- differential expressed in IUGR-Reactome pathway**

| #term ID | term description | observed g | backgroun | FDR | Protein name |
|---|---|---|---|---|---|
| HSA-2214320 | Anchoring fibril formation | 2 | 15 | 0.00064 | COL4A1,COL4A2 |
| HSA-2243919 | Crosslinking of collagen fibrils | 2 | 18 | 0.00064 | COL4A1,COL4A2 |
| HSA-3000157 | Laminin interactions | 2 | 30 | 0.00064 | COL4A1,COL4A2 |
| HSA-3000480 | Scavenging by Class A Receptors | 2 | 18 | 0.00064 | COL4A1,COL4A2 |
| HSA-140877 | Formation of Fibrin Clot (Clotting Cascade) | 2 | 39 | 0.00077 | SERPINE2,TFPI |
| HSA-419037 | NCAM1 interactions | 2 | 42 | 0.00077 | COL4A1,COL4A2 |
| HSA-8948216 | Collagen chain trimerization | 2 | 44 | 0.00077 | COL4A1,COL4A2 |
| HSA-1442490 | Collagen degradation | 2 | 64 | 0.00083 | COL4A1,COL4A2 |
| HSA-186797 | Signaling by PDGF | 2 | 55 | 0.00083 | COL4A1,COL4A2 |
| HSA-3000171 | Non-integrin membrane-ECM interactions | 2 | 58 | 0.00083 | COL4A1,COL4A2 |
| HSA-3000178 | ECM proteoglycans | 2 | 75 | 0.00091 | COL4A1,COL4A2 |
| HSA-216083 | Integrin cell surface interactions | 2 | 83 | 0.00099 | COL4A1,COL4A2 |
| HSA-9006934 | Signaling by Receptor Tyrosine Kinases | 3 | 437 | 0.00099 | CDH5,COL4A1,COL4A2 |

**S3 A: The 33 transcription factors (TFs) regulating the expression of genes encoding proteins in the placental secretome map with indication of the target genes differentially expressed in the placenta in pregnancy complications.**

| TF | TF targets from the placental secretome | TF targets from the placental secretome that are altered in the placenta in pregnancy complications (reported in Figure 4A) |
|---|---|---|
| **ARNT2** | APOE | In PE: GPX3, LIFR, POSTN |
| | BAG6 | In PE, SGA: IGFBP4 |
| | GPX3 | In GDM: ITGB3 |
| | IGFBP4 | |
| | ITGB3 | |
| | LAMA2 | |
| | LIFR | |
| | LTF | |
| | PFN1 | |
| | POSTN | |
| **ELF3** | FN1 | In IUGR, PE, GDM: FN1 |
| | TIMP3 | In GDM, PE, SGA, IUGR: TIMP3 |
| **PLAG1** | IGF2 | In GDM, PE, SGA, IUGR: IGF2 |
| | QSOX1 | |
| **SP2** | SERPINE1 | In GDM, PE: SERPINE1 |
| | SERPINH1 | |
| MEF2D | COL1A2 | In PE: COL1A2, SOD1, TGFB2 |
| | COL3A1 | |
| | SOD1 | |
| | TGFB2 | |
| **IRF3** | ANXA4 | In GDM, PE, LGA: ANXA4 |
| | B2M | In GDM, PE: CCL2 |
| | CCL2 | In GDM, PE, IUGR: FN1 |
| | FN1 | |
| | ISG15 | |
| | PNP | |
| **CREB1** | FLT1 | In GDM, PE, SGA, IUGR: FLT1 |
| | FN1 | In GDM, PE, IUGR: FN1 |
| **FOS** | CLU | In GDM, PE: HSPA5 |
| | COL6A1 | In PE: SERPINB9, TIMP1 |
| | FETUB | |
| | GKN1 | |
| | HSPA5 | |
| | SERPINB9 | |
| | TIMP1 | |
| **MYCN** | ACTG1 | In PE: ACTG1 |
| | FN1 | In GDM, PE, IUGR: FN1 |
| **NFYC** | COL1A1 | In GDM, PE: COL1A1 |
| | FTH1 | |
| **EHF** | ANPEP | In PE: S100A9, TIMP1 |
| | S100A9 | In GDM, PE: SERPINE1 |
| | SERPINE1 | In GDM, PE, IUGR: TFPI |
| | TFPI | |
| | TIMP1 | |
| **SPA1** | ADA | In PE: ADA, COL1A2, CTSD, SOD1, TFPI2, TIMP1 |
| | AEBP1 | In GDM, PE: CCL2, COL1A1, SERPINE1 |
| | APOE | In GDM, PE, SGA, IUGR: FLT1, IGF2, TIMP3 |
| | CAT | In GDM, PE, IUGR: FN1 |
| | CCL2 | In PE, SGA: IGFBP4, MMP2 |
| | COL1A1 | In GDM: ITGB3, SPARC |
| | COL1A2 | |
| | CTSD | In IUGR: SERPINE2 |
| | FLT1 | |
| | FN1 | |
| | IGF2 | |
| | IGFBP4 | |
| | ITGB3 | |
| | LAMA1 | |
| | MMP2 | |
| | PGK1 | |
| | PKM | |
| | PROCR | |
| | SCT | |
| | SERPINE1 | |
| | SERPINE2 | |
| | SOD1 | |
| | SPARC | |
| | TFPI2 | |
| | TIMP1 | |
| | TIMP3 | |
| | TNC | |
| **KLF3** | AHNAK | In IUGR, PE: ANXA5 |
| | ANXA5 | |
| | ANXA6 | |

| | | |
|---|---|---|
| | APOE | |
| | CAPG | |
| | PCYOX1 | |
| | PRKACA | |
| | SERPINH1 | |
| **ETV5** | FN1 | In GDM, PE, IUGR: FN1 |
| | ITGB1 | In GDM, PE: ITGB1 |
| | KRT13 | In PE, SGA: MMP2 |
| | MMP2 | In PE: S100A6 |
| | S100A6 | In GDM, PE, SGA, IUGR: TIMP3 |
| | TIMP3 | |
| **KLF4** | ALB | In GDM, PE, SGA, IUGR: FLT1 |
| | ARG1 | In GDM, PE, IUGR: FN1, SEMA3F |
| | FLT1 | In PE, IUGR: INHBA |
| | FN1 | In GDM, PE: SERPINE1 |
| | INHBA | In PE: TGFB2 |
| | KRT13 | |
| | LAMA1 | |
| | SEMA3F | |
| | SERPINE1 | |
| | SERPINH1 | |
| | TGFB2 | |
| **NRF1** | CD47 | |
| **FOXO3** | ANGPT2 | In GDM, PE, SGA, IUGR: ANGPT2 |
| | COL4A1 | In IUGR: COL4A1 |
| | IGFBP7 | In PE, IUGR: INHBA |
| | INHBA | In GDM, PE, SGA: NAMPT |
| | NAMPT | In PE: PRCP, TGFB2 |
| | PRCP | In GDM, PE: SERPINE1 |
| | SERPINE1 | |
| | TGFB2 | |
| **FOXO4** | NAMPT | In GDM, PE, SGA: NAMPT |
| | SERPINE1 | In GDM, PE: SERPINE1 |
| **ATF3** | GCG | In GDM: HSPA5 |
| | HSPA5 | |
| **SMAD2** | BGN | In LGA, PE, SGA, IUGR: BGN |
| | COL1A2 | In PE: COL1A2 |
| | ITGB1 | In GDM, PE: ITGB1 |
| | SERPINE1 | In GDM, PE: SERPINE1 |
| **TBP** | COL1A1 | In IUGR: COL4A1 |
| | FN1 | In GDM, PE, IUGR: FN1 |
| | IGF2 | In GDM, PE, SGA, IUGR: IGF2 |
| **MAX** | YBX1 | |
| **MECP2** | ANXA1 | In PE: ANXA1, LGALS1, TGFB2 |
| | HSPA5 | In GDM: HSPA5 |
| | IGF2 | In GDM, PE, SGA, IUGR: IGF2 |
| | LGALS1 | |
| | TGFB2 | |
| **SP3** | AEBP1 | In GDM, PE, SGA, IUGR: ANGPT2, FLT1, IGF2 |
| | ANGPT2 | In GDM, PE: CCL2, COL1A1, SERPINE1 |
| | CCL2 | In PE: COL1A2, PKM, TIMP1 |
| | COL1A1 | In PE, SGA: MMP2 |
| | COL1A2 | In GDM, PE, IUGR: PLAU |
| | FLT1 | In GDM: SPARC |
| | IGF2 | |
| | LAMA1 | |
| | MMP2 | |
| | PGK1 | |
| | PKM | |
| | PLAU | |
| | PROCR | |
| | SERPINE1 | |
| | SERPINH1 | |
| | SPARC | |
| | TIMP1 | |
| **NFYA** | CALR | In GDM: CALR, HSPA5 |
| | CAT | In PE: COL1A2, PKM |
| | COL1A2 | In GDM, PE: HSPA5 |
| | FTH1 | |
| | HSPA5 | |
| | PKM | |
| **SMAD4** | ANGPT2 | In GDM, PE, SGA, IUGR: ANGPT2, TIMP3 |
| | APOA1 | In LGA, PE, SGA, IUGR: BGN |
| | BGN | In GDM, PE: CCL2, SERPINE1 |
| | CCL2 | In PE: COL1A2, TGFB2, TIMP1 |
| | COL1A2 | In GDM, PE: ITGB1 |
| | CSF1 | In GDM, PE, IUGR: PLAU |
| | ITGB1 | In GDM, PE, SGA: THBS1 |
| | PLAU | |
| | SERPINE1 | |

| | | |
|---|---|---|
| | TGFB2 | |
| | THBS1 | |
| | TIMP1 | |
| | TIMP3 | |
| | TNC | |
| **ARNT** | CTSD | In PE: CTSD, TFRC |
| | TFRC | |
| **HNF1B** | AFP | In PE GDM, LGA: ANXA4 |
| | ALB | In GDM, PE, IUGR: PLAU |
| | ANXA4 | In GDM: RNASE4, SPARC |
| | LUM | In PE, IUGS, SGA: TTR |
| | PLAU | |
| | RNASE4 | |
| | SPARC | |
| | TTR | |
| **KLF6** | ARG1 | In GDM, PE: CCL2, COL1A1 |
| | CCL2 | |
| | COL1A1 | |
| | MSLN | |
| | SERPINH1 | |
| **FOS** | CLU | In GDM, PE: HSPA5 |
| | COL6A1 | In PE: TIMP1 |
| | FETUB | |
| | GKN1 | |
| | HSPA5 | |
| | SERPINB9 | |
| | TIMP1 | |
| **KLF15** | FABP3 | |
| | FABP5 | |
| **CREB1** | FLT1 | In GDM, PE, SGA, IUGR: FLT1 |
| | FN1 | In GDM, PE, IUGR: FN1 |
| **USF2** | IGF2R | |
| **IRF2** | ARG1 | In GDM: CTSS |
| | B2M | |
| | C1QTNF1 | |
| | CHIL3 | |
| | CLIC1 | |
| | CTSS | |
| | ISG15 | |
| **PRDM1** | DCN | In PE: DCN |
| | SERPINE1 | In GDM, PE: SERPINE1 |
